# From the scanner to daily life: Neural expression of the Picture-Induced Negative Emotion Signature is associated with real-world daily and momentary negative affect in midlife adults

**DOI:** 10.64898/2026.06.03.729931

**Authors:** E. Lydia Wu-Chung, William D. Eckerle, Anna L. Marsland, Thomas E. Kraynak, Thomas W. Kamarck, Peter J. Gianaros

## Abstract

Functional magnetic resonance imaging (fMRI) is widely used to map brain systems that process affective stimuli and predict concurrent affective experiences in the scanner. However, the extent to which affect-related fMRI activity in the scanner predicts affective experiences in daily life remains unclear. This study examined whether a multivariate brain pattern previously validated to predict negative affect to emotional images during fMRI testing (the Picture-Induced Negative Emotion Signature; PINES) associates with daily negative affect measured over 7-10 days via daily diaries and ecological momentary assessment. Among 287 midlife adults, those exhibiting greater PINES pattern expression showed greater daily (end-of-day) and hourly negative affect. These findings were not explained by demographic factors and trait negative affect. A cross-validated whole-brain pattern of negative affect generalizes beyond the fMRI testing context to predict negative affect severity in daily life.

---

Task-based functional MRI (fMRI) has been foundational for studying the neurobiology of emotion. Through carefully controlled testing paradigms, various aspects of emotion processing and regulation have been attributed to activity in distributed brain systems. Although testing paradigms administered during fMRI rarely mirror the complexity of real-world experiences, fMRI findings are usually interpreted as corresponding to psychological processes occurring outside the testing context. However, few studies investigate associations between task-evoked fMRI activity and real-world psychological experiences. Given increasing interest in understanding the neural bases of negative affect and affective disorders, determining whether neural patterns evoked by task fMRI meaningfully relate to daily emotional experiences is important.

Previous studies report significant associations between task-elicited brain activity and socioemotional patterns in everyday life. In a sample of 42 young adults, metabolic activity in brain regions engaged by a social rejection task (e.g., amygdala, periaqueductal gray, dorsal anterior cingulate cortex) correlated positively with momentary social distress in daily life.

Among 208 subjects, Hur et al. (2022) examined associations between an anxiety-provoking fMRI task and negative affect patterns in daily life. Activation in frontocortical (e.g., dorsolateral prefrontal cortex, midcingulate cortex, anterior insula, frontal operculum), but not subcortical regions (e.g., amygdala, periaqueductal gray, bed nucleus of the stria terminalis), correlated with stress-induced negative affect in daily life (Hur et al., 2022). While these studies provide some support for the ecological validity and generalizability of task-based socioemotional fMRI tasks, they rely on mass univariate correlational approaches or region-of interest (ROI) analysis, which have problematic psychometric properties and poor replicability (Elliott et al., 2020). Because replication remains a significant challenge among fMRI studies with small sample sizes and those using mass univariate or ROI analytic approaches (Elliott et al., 2020; Turner et al., 2020), studies focused on more reproducible brain patterns and daily psychological processes are needed (Kragel et al., 2021).

Multivariate methods using machine learning and cross-validation are alternative approaches to mass univariate methods for characterizing replicable and distributed patterns of coordinated brain activity. Based on the assumption that psychological processes rely on many interactive brain systems rather than single brain regions operating in isolation, multivariate approaches capitalize on the high-dimensional nature of correlated fMRI data to identify distributed activation patterns or whole-brain signatures (Woo et al., 2017). A common approach is to employ machine-learning methods to derive a whole-brain pattern from a training sample that is subsequently cross-validated by predicting out-of-sample observations. Multivariate neuroimaging approaches increase test-retest reliability (Kragel et al., 2021) and, therefore, the potential for replication across fMRI studies (Noble et al., 2021). To our knowledge, no study has examined whether pre-established multivariate brain patterns associated with emotional processing relate to emotional experiences in daily life.

A multivariate brain pattern relevant to the study of daily affective experiences is the Picture-Induced Negative Emotion Signature (PINES) developed by Chang et al. (2015). This whole-brain pattern was trained from fMRI data to predict in-scanner negative affect ratings to negative and neutral images. In other words, the PINES reflects a brain-based model of self-reported negative affect intensity during fMRI. Model weights in the PINES map correspond to regions implicated in emotion processing (Lindquist et al., 2012). Given the specificity of the PINES to state negative affect, a pertinent and unexplored question is whether individual differences in PINES expression predicts negative affect in daily life.

Examining the association between neural measures of negative affect and everyday negative affect can be achieved through daily diary measures and ecological momentary assessment (EMA; Hufford et al., 2001; Kamarck et al., 2011). These involve assessing negative affect at the end of each day (daily diary) or at various moments throughout the day (EMA), wherein individuals report on their current affect in the places and at the times of their typical activities (e.g. at work or home). These methods enhance the reliability of measurement by removing or reducing recall bias, as affect ratings are provided in near real time or after a short recall period. These attributes of EMA and daily diary assessments make them ideal for establishing the external validity of neural measures of affective processes, particularly the PINES.

By integrating task fMRI, EMA and daily diary approaches, the present study examined whether individual differences in whole-brain patterns of negative affect expression encoded by the PINES relate to negative affect in daily life. We hypothesized that individuals showing brain patterns more similar to the PINES (i.e., greater PINES expression) would also report more negative affect in daily life. Some studies have reported differences between end-of-day vs. momentary reports of emotion and task-evoked brain activity (Eisenberger et al., 2007). Accordingly, we examined whether PINES expression was associated with end-of-day and hourly measures.

## Method

### Research Transparency Statement

Conflicts of interest: None of the authors have conflicts of interest to disclose. Preregistration: Hypotheses and methods were not preregistered. Materials, Data, and Analysis scripts: Some study materials, de-identified data, and analysis code are publicly available (https://osf.io/pnjuv/overview?view_only=b054e1d21ce742a3af54e88b0f196eba). Funding: This research was supported by grants from the National Heart, Lung, and Blood Institute (R01-HL089850, P01-HL040962, F32HL176202) and National Institute of Diabetes and Digestive and Kidney Diseases (R01-DK110041). Artificial intelligence: Artificial intelligence (Arena) assisted in the development of analytic code for mixed level modeling and figures; it was not used to generate any data or text for the writing of this manuscript. AI-generated code was reviewed by authors to ensure accuracy.

### Participants

Participants were volunteers of the Neurobiology of Adult Health (NOAH) project, which was designed to examine the neural correlates of heart disease risk in midlife. A total of 366 participants were enrolled between Feb 2019 and October 2021 (Mean age: 42.09, SD = 8.65; 63% female, 83% White, 8% Black, 6% Asian). To be eligible for the study, participants had to be English speaking and aged 28-56. Participants were excluded if they were actively taking psychotropic or immune-related medication, diagnosed with chronic diseases, had high blood pressure, or had barriers to completing electronic diary or ambulatory assessments (see supplemental material for a detailed list).

### Study design

Participants completed visits during a baseline phase and a 2-year follow-up phase. Due to the wide range of biobehavioral assessments incorporated into the parent study, data at each time point was acquired across several visits over a span of 2-3 months. This study utilized self-report questionnaires, ecological momentary assessments, daily dairy measures, and functional magnetic resonance imaging (fMRI) data from the baseline visit. Daily diary assessments were acquired in the week following self-report questionnaire data collection and before the fMRI visit (Average: 43.8 days, SD = 47.45 days).

After the baseline visit, participants underwent approximately 7-10 days of daily life monitoring for EMA and daily diary data collection. This included four “full monitoring” days and approximately six “partial monitoring” days. During the four “full monitoring” days, participants completed hourly surveys and end of day surveys, both of which included measures of negative affect (see “Measures” below). In contrast to full monitoring days, “partial monitoring” days consisted of only end of day surveys. To ensure understanding of procedures, participants underwent an initial practice day, during which they completed all procedures of a full monitoring day and received a brief phone call from study staff to check-in and provide feedback. Participants faced with unexpected challenges, technical issues, or misunderstandings underwent extra monitoring days, resulting in some individuals with more than 10 days of data.

For the present study, only participants who completed at least one assessment of daily affect and the affective fMRI task were included. Of the 366 participants enrolled in NOAH, 299 and 320 completed the affective fMRI task or daily diary, respectively. After excluding participants who had unusable neuroimaging data (n=4) and those who didn’t have both fMRI and daily diary data (n=32), 288 participants remained. We further excluded one participant from the final analysis due to being the sole representative of their sex category (i.e., intersex, transgender), which prevented meaningful statistical comparison. Thus, the final analytic sample consisted of 287 subjects.

## Measures

### Self-reported negative affect

#### Trait negative affect

At the baseline visit, the negative affect subscale of the 20-item Positive and Negative Affect Scale (PANAS) (Watson et al., 1988) was used to assess trait negative affect. The following items were part of the negative affect subscale: ‘distressed’, ‘upset’, ‘guilty’, ‘hostile’, ‘scared’, ‘irritable’, ‘ashamed’, ‘nervous’, ‘jittery’, ‘afraid.’ For each item, participants rated how they felt <u>in general</u>, using a 5-point Likert scale: 1 = very slightly or not at all; 2 = a little; 3 = moderately; 4 = quite a bit; 5 = extremely. Higher values indicate more negative affect. Internal consistency for the negative affect subscale was good (Cronbach’s ⍺ = .85). Positive affect (Cronbach’s ⍺ = .87) was not of primary interest; however, post-hoc analyses with positive affect were explored to examine robustness of study findings.

### EMA and Daily Diary Measures

The NOAH study involved two forms of EMA data collection for negative affect ratings, hourly surveys and end-of-day surveys. Hourly surveys consisted of a revised version of the Diary of Ambulatory Behavioral States (DABS; Kamarck et al., 1998), a 46-item self-report measure designed for repeated real-time assessment of daily experiences. Multi-item scales assess affect, activity, and social interactions, as well as common behavioral factors of interest to clinical and health psychology research (e.g., physical activity, eating, etc).

Daily and hourly negative affect was measured using 3 items on the DABS: sad, frustrated/angry, and nervous/tense. For hourly assessments, participants rated how they felt with respect to each of these states just before they began the survey, on a scale from 0 being “No” to 10 being “Yes”. For end-of-day negative affect, participants responded to the prompt: *“In general how did you feel today…”* using the same scale. Ratings on these three variables were averaged together to create a composite, negative affect score. Given the limited temporal scope of both hourly and end-of-day based measures, we conceptualized these as measures of state, rather than trait, negative affect.

### Whole-brain pattern of negative affect

#### IAPS task

Participants completed an affective viewing and response task, identical to previously described protocols (Gianaros et al., 2014; Ochsner et al., 2002, 2004). Briefly, participants viewed 30 unpleasant images and 15 neutral images from the International Affective Picture System (IAPS; see supplemental material for IAPS IDs)(Lang et al., 2008). Images were preceded by a 2-s cue indicating whether participants should ‘Look’ and attend to images normally or ‘Decrease’ and alter how they thought about the image. Following a 7-s IAPS image presentation, participants rated their emotional state (“How negative do you feel?”) using a 5-point Likert scale (1 = neutral, 5 = strongly negative) during a 4-s response period. A variable 1-3s rest period followed each rating period. Participants were first trained prior to completing the task in the scanner. The entire task lasted 11 min and 16 s (15 ‘Look Neutral’ trials, 15 ‘Look Negative’ trials, 15 ‘Decrease Negative’ trials).

### Magnetic Resonance Imaging Data Acquisition

Imaging was conducted on a 3T PRISMA scanner (Siemens) with a 64-channel head coil. A T1-weighted MPRAGE structural image was obtained using the following parameters: 176 sagittal slices (1mm thick, no gap, matrix size = 176×176 voxels [FOV = 256×256mm]); TR = 2300ms; inversion time = 900ms; TE = 1.99ms; and FA = 9°. Functional BOLD data acquisition parameters were as follows: matrix size = 106 × 106 voxels (FOV = 212 × 212 mm), TR = 2000 ms, TE = 30 ms, and FA = 79°. 69 slices per volume were collected along an anterior-to-posterior encoding direction. Each volume was 2 mm in thickness, with no gap (280 task volumes in total, excluding 4 discarded volumes).

### MRI Preprocessing

MRI data were preprocessed using statistical parametric mapping software (SPM12; https://www.fil.ion.ucl.ac.uk/spm/). T1-weighted MPRAGE images were classified into 6 tissue types for spatial preprocessing, followed by computation of biased-corrected and deformation field maps. Slice-timing correction was applied to account for variations in acquisition time. Realignment of functional images to the first image of the series was then performed via 6-parameter rigid-body transformation, using the re-slice step to match the first image on a voxel-by-voxel basis. Realigned images were coregistered to each subject’s bias-corrected and skull-stripped MPRAGE image. Coregistered images were normalized to Montreal Neurological Institute (MNI) space and subsequently smoothed with a 6-mm full-width-at-half-maximum (FWHM) Gaussian kernel. Preprocessed images were manually inspected to ensure no errors occurred that would render images unusable for analysis.

For primary analyses, first-level general linear models (GLMs) were constructed for each subject to derive subject-level contrast maps subsequently used to compute PINES expression scores. For the GLM, task events were modeled by rectangular waveforms convolved with the default hemodynamic response function in SPM12. Regressors modeled trial events (i.e., cue, IAPS scene, rating period, rest). Six realignment parameters from preprocessing were included in GLMs as nuisance regressors, and low frequency artifacts were identified and removed using a high-pass filter (128 s). Error variance attributed to noise and artifacts in fMRI time series was estimated and subsequently weighted by restricted maximum likelihood estimation, using robust weighted least squares (WLS) toolbox, v4.0 (Diedrichsen & Shadmehr, 2005). Linear contrasts of different trial conditions were computed. For this study, only the ‘Look Negative vs. Look Neutral’ contrast was utilized as these trials were used in the development and validation of the PINES.

### PINES expression scores

The Spearman’s rank correlation coefficient between unthresholded Picture Induced Negative Emotion Signature (PINES) – which had been originally trained and cross-validated to predict subjective ratings of negative affect (Chang et al., 2015) – and the ‘Look Negative vs. Look Neutral’ activation map for each subject was computed to derive a composite summary measure (pattern expression) reflecting the degree of similarity to the PINES. As a result, higher expression scores indicate greater overall similarity to the PINES or a whole-brain pattern indicative of more negative affect.

### Covariates

Age, sex, and years of education were included as covariates in the model. Sex was dummy coded (0 = male, 1 = female).

### Analytic approach

#### Primary analyses

All analyses were conducted in R Studio. Preliminary model testing revealed that 46% of the variance in daily negative affect was attributed to within-person variation (ICC: .54). Thus, linear mixed-effects models fit by restricted maximum likelihood (REML) were used to test study hypotheses. The -2 log-likelihood value, Akaike Information Criterion (AIC), and Bayesian Information Criterion (BIC) were used to determine whether fixed, random, and/or polynomial effects should be incorporated into the model. For analyses of daily negative affect, the best-fitting model incorporated a random intercept for subject, which allows between-person differences in overall negative affect. For analyses of hourly negative affect, the best-fitting model included linear and quadratic terms of hour of the day (military time, person mean-centered) as fixed effects and subject and day nested within subject as random intercepts. The quadratic term allowed for curvilinear diurnal patterns in negative affect and a random intercept for day nested within person allowed for day-specific fluctuations in negative affect. Negative affect was regressed on the main variable of interest (e.g., PINES expression score) in unadjusted and adjusted models. Unadjusted models only included PINES expression score and the above-mentioned fixed and random effects as predictors. Adjusted models further included trait negative affect, age, sex, and education as covariates. Due to three participants missing trait negative affect data, sample sizes for unadjusted and adjusted models were 287 and 284, respectively.

For completeness of reporting, we also analyzed PINES expression scores computed by the alternative dot product method (a vector-based indicator of alignment between the PINES and each subject’s contrast map). The correlation between Spearman-based and dot-product scores was high in this sample (*r* = .80; see Supplemental Material). As reported below and in the Supplemental Material, the minimally-adjusted associations between hourly (0.12 vs. 0.10) and end-of-day (0.21 and 0.14) negative affect with both expression scores were comparably small with respect to the magnitude of their effect sizes. Findings for dot product analyses can be found in the Supplemental Material.

### Ancillary analyses

We conducted several ancillary analyses to verify the validity and robustness of our findings: **1) Outlier detection and removal of practice days.** Cook’s Distance was used to identify the presence of influential outliers; subjects with values exceeding 1 were excluded in sensitivity analyses. We also examined whether observed effects were affected by the inclusion of practice days by rerunning momentary analyses and excluding data from practice days; **2) Validation of the PINES in the NOAH cohort.** To our knowledge, no study has validated the PINES model in a separate cohort using the same IAPS task. Thus, we reconstructed GLMs for each subject in accordance with the original PINES modeling approach (see Supplement) and examined how well the PINES predicted within-subject, negative affect ratings of Look Negative and Look Neutral trials. The ecological validity of laboratory emotion paradigms is also not commonly explored. Thus, to determine whether negative affect ratings in the scanner were associated with everyday negative affect patterns, we examined the association between-subject negative affect ratings of Look Negative vs. Look Neutral images and negative affect in daily life; **3) Positive affect.** To examine whether observed patterns held after accounting for positive affect, we controlled for trait and state (end-of-day) positive affect. We also explored whether PINES expression scores were sensitive to emotional valence (negative or positive affect) by examining whether PINES expression scores predicted daily and momentary positive affect; **4) Amygdalar contributions.** Amygdala activity is frequently implicated in the experience of negative affect. To examine whether negative affect in daily life could be explained solely by mean amygdala activation to Look Negative vs. Look Neutral images, we reran analyses using a) a bilateral anatomical mask of the amygdala and b) a PINES map lesioned for the amygdala.

## Results

Descriptive sample information is detailed in Table 1. Bivariate correlations (Table 2) revealed that PINES expression scores were positively correlated with mean end-of-day negative affect (*p* = .024) but not trait (*p* = .761) or mean hourly negative affect (*p* = .105). Between-subject in-scanner negative affect ratings also correlated positively with end-of-day negative affect (*p* = .043) but not trait (*p* = .694) or mean hourly negative affect (*p* = .135). End-of-day negative affect correlated with all trait and state measures of positive and negative affect (*p*’s < .001). For the current analytic sample, 95% of the participants completed 4 or more full monitoring days.

**Table 1.**
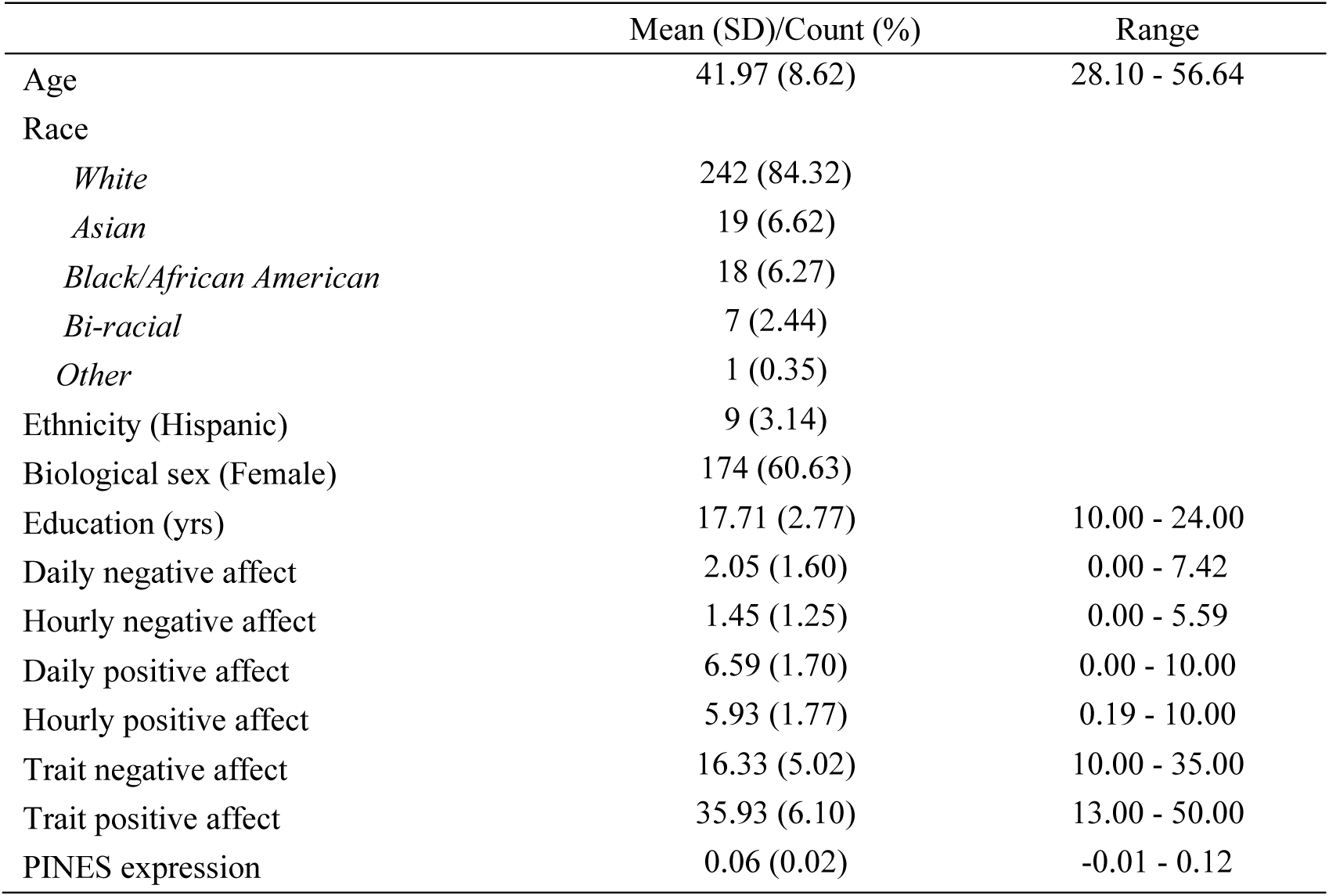
Descriptive characteristics of analytic sample (N=287)

|  | Mean (SD)/Count (%) | Range |
| --- | --- | --- |
| Age | 41.97 (8.62) | 28.10 - 56.64 |
| Race |  |  |
| *White* | 242 (84.32) |  |
| *Asian* | 19 (6.62) |  |
| *Black/African American* | 18 (6.27) |  |
| *Bi-racial* | 7 (2.44) |  |
| *Other* | 1 (0.35) |  |
| Ethnicity (Hispanic) | 9 (3.14) |  |
| Biological sex (Female) | 174 (60.63) |  |
| Education (yrs) | 17.71 (2.77) | 10.00 - 24.00 |
| Daily negative affect | 2.05 (1.60) | 0.00 - 7.42 |
| Hourly negative affect | 1.45 (1.25) | 0.00 - 5.59 |
| Daily positive affect | 6.59 (1.70) | 0.00 - 10.00 |
| Hourly positive affect | 5.93 (1.77) | 0.19 - 10.00 |
| Trait negative affect | 16.33 (5.02) | 10.00 - 35.00 |
| Trait positive affect | 35.93 (6.10) | 13.00 - 50.00 |
| PINES expression | 0.06 (0.02) | -0.01 - 0.12 |

**Table 2.** Bivariate Pearson correlations of main variables of interest.

| Variable | 1 | 2 | 3 | 4 | 5 | 6 | 7 | 8 | 9 |
| --- | --- | --- | --- | --- | --- | --- | --- | --- | --- |
| 1. PINES expression |  |  |  |  |  |  |  |  |  |
| 2. Daily NA | .13* |  |  |  |  |  |  |  |  |
| 3. Hourly NA | .10 | .88** |  |  |  |  |  |  |  |
| 4. Trait NA | .02 | .54** | .50** |  |  |  |  |  |  |
| 5. Daily PA | -.06 | -.52** | -.48** | -.32** |  |  |  |  |  |
| 6. Hourly PA | -.05 | -.41** | -.40** | -.28** | .88** |  |  |  |  |
| 7. Trait PA | .02 | -.29** | -.23** | -.28** | .51** | .49** |  |  |  |
| 8. Age (yrs) | .03 | -.15* | -.10 | -.12* | .08 | .10 | .11 |  |  |
| 9. Education (yrs) | -.03 | .10 | .09 | -.01 | -.11 | -.13* | -.03 | -.17** |  |
| 10. NA rating (Look Neg v. Look Neu) | .28** | .12* | .09 | -.02 | .09 | .04 | .12 | .02 | .06 |

*Note.* \* indicates $p < .05$ . \*\* indicates $p < .01$ . NA = negative affect. Daily affect was based on end-of-day reports of affect. Hourly affect is the within-person average of affect acquired across all hourly data. NA rating is the between-subject negative affect ratings toward IAPS Look Negative vs Look Neutral images.

The average number of days spent in full monitoring was 5.51 days (range, 2-9). On average, participants completed 9.09 daily surveys (range, 4-16) and 10.28 hourly surveys per day (range, 1-19).

### Daily negative affect

Detailed statistical results from linear mixed models of daily (end-of-day) negative affect regressed on variables of interest are displayed in Table 3. In unadjusted models, PINES expression scores positively associated with end-of-day negative affect (*p* = .025). These patterns remained statistically significant after controlling for demographic covariates and trait negative affect (*p* = .007, see Figure 1). In ancillary tests, PINES expression scores remained statistically predictive of daily end-of-day negative affect after additionally controlling for trait and end-of-day positive affect (*p* = .009). No influential outliers were detected. These patterns were specific to negative affect, as PINES expression scores were not predictive of daily positive affect (*p* = .203).

**Figure 1.**
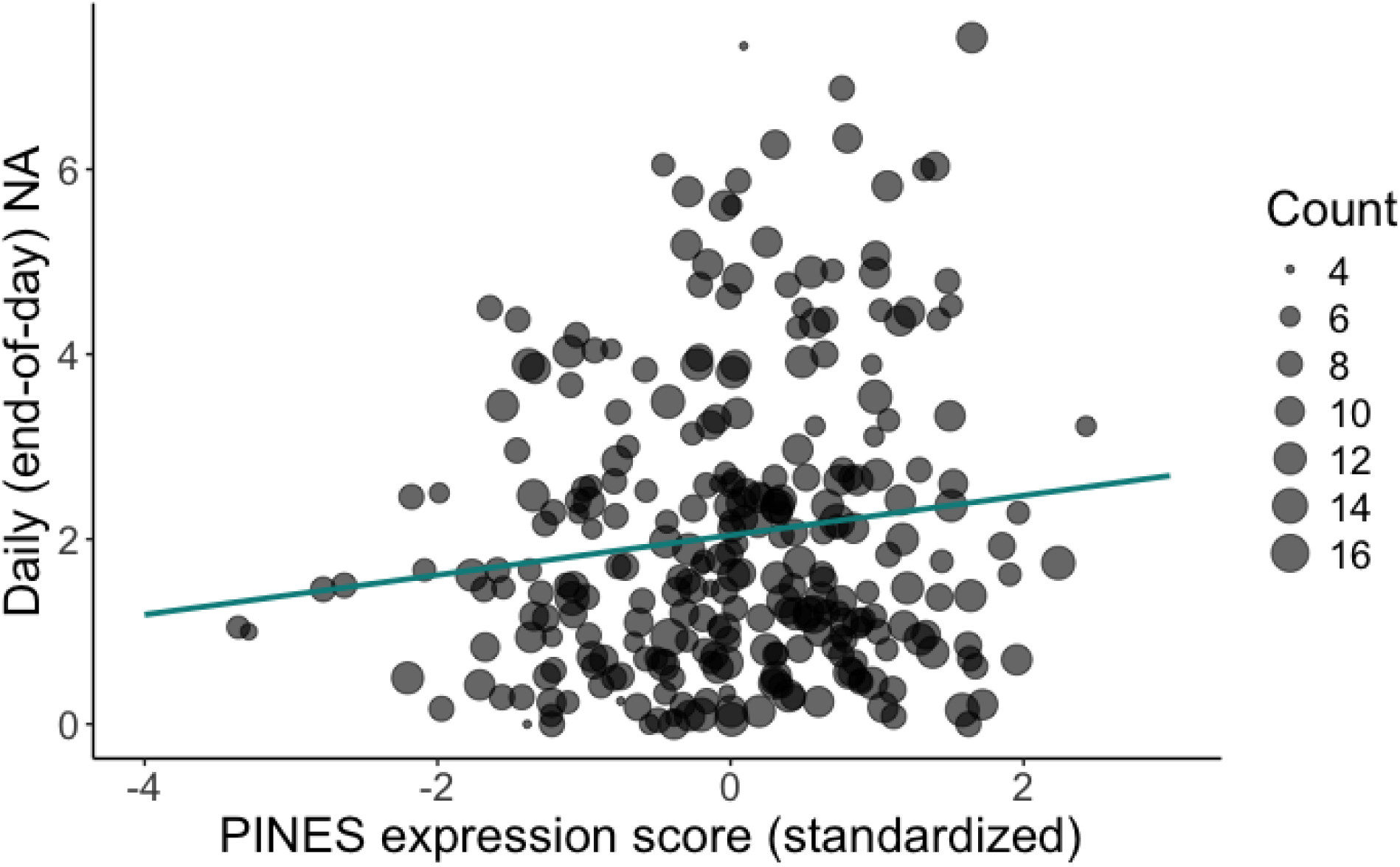
The association between PINES expression scores and daily negative affect (NA). *Note:* This reflects the model that controlled for age, sex, education, and trait negative affect. For graphical purposes, individual data points were aggregated into a mean for each subject, such that each subject is represented by a dot and the size of the dot represents the number (count) of diary entries.

**Table 3.**
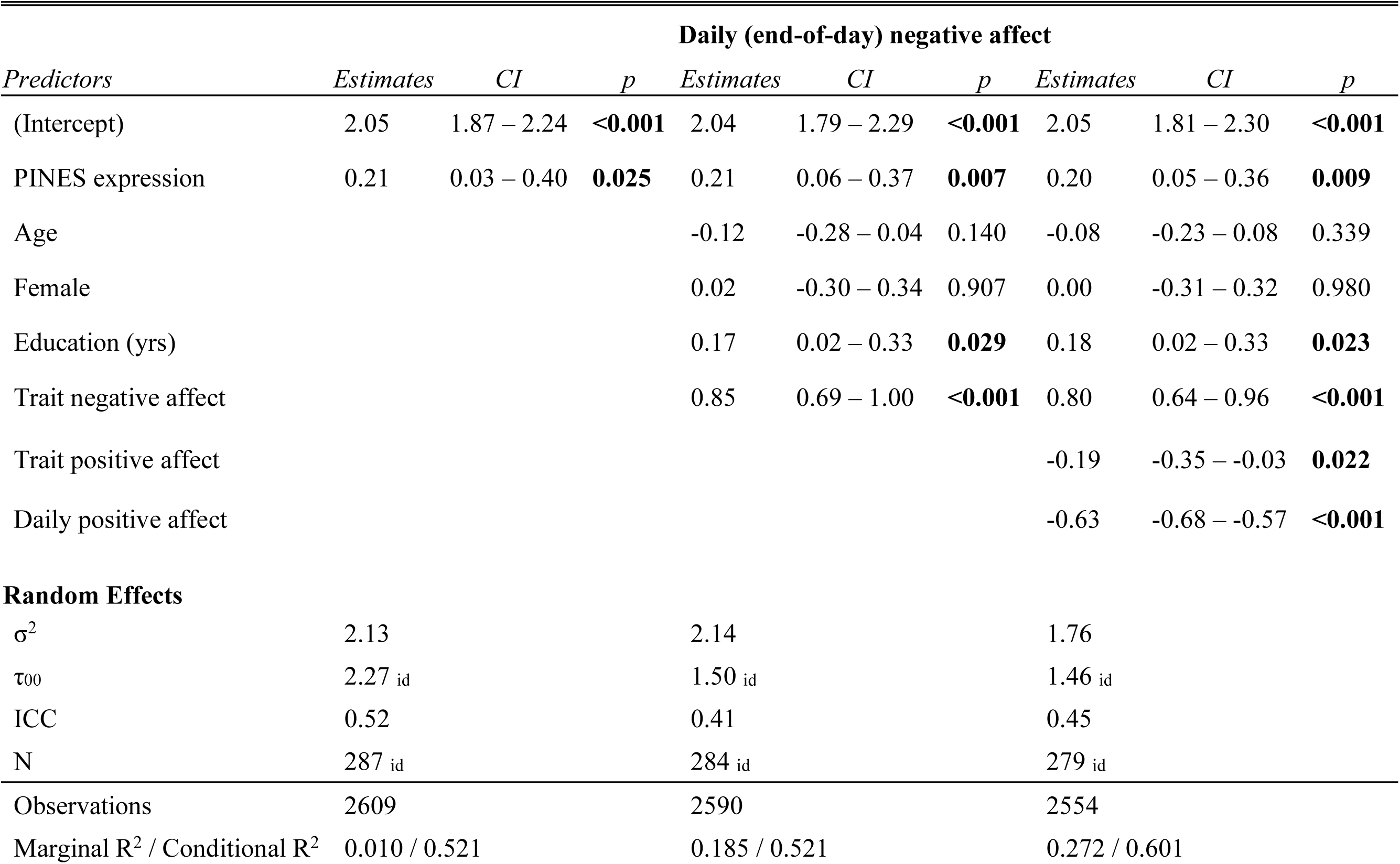
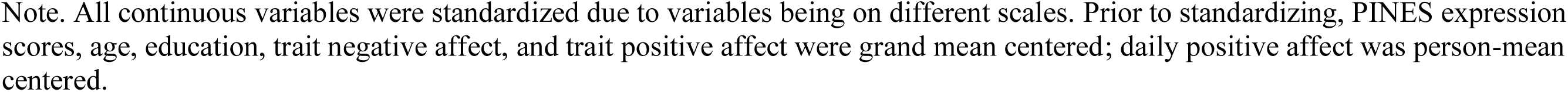
Linear mixed model results regressing daily negative affect on PINES expression scores and covariates.

| *Predictors* | **Daily (end-of-day) negative affect** |  |  |  |  |  |  |  |  |
| --- | --- | --- | --- | --- | --- | --- | --- | --- | --- |
|  | *Estimates* | *CI* | *p* | *Estimates* | *CI* | *p* | *Estimates* | *CI* | *p* |
| (Intercept) | 2.05 | 1.87 – 2.24 | **<0.001** | 2.04 | 1.79 – 2.29 | **<0.001** | 2.05 | 1.81 – 2.30 | **<0.001** |
| PINES expression | 0.21 | 0.03 – 0.40 | **0.025** | 0.21 | 0.06 – 0.37 | **0.007** | 0.20 | 0.05 – 0.36 | **0.009** |
| Age |  |  |  | -0.12 | -0.28 – 0.04 | 0.140 | -0.08 | -0.23 – 0.08 | 0.339 |
| Female |  |  |  | 0.02 | -0.30 – 0.34 | 0.907 | 0.00 | -0.31 – 0.32 | 0.980 |
| Education (yrs) |  |  |  | 0.17 | 0.02 – 0.33 | **0.029** | 0.18 | 0.02 – 0.33 | **0.023** |
| Trait negative affect |  |  |  | 0.85 | 0.69 – 1.00 | **<0.001** | 0.80 | 0.64 – 0.96 | **<0.001** |
| Trait positive affect |  |  |  |  |  |  | -0.19 | -0.35 – -0.03 | **0.022** |
| Daily positive affect |  |  |  |  |  |  | -0.63 | -0.68 – -0.57 | **<0.001** |
| **Random Effects** |  |  |  |  |  |  |  |  |  |
| $\sigma^2$ | 2.13 | | | 2.14 | | | 1.76 | | |
| $\tau_{00}$ | 2.27<sub>id</sub> | | | 1.50<sub>id</sub> | | | 1.46<sub>id</sub> | | |
| ICC | 0.52 |  |  | 0.41 |  |  | 0.45 |  |  |
| N | 287<sub>id</sub> |  |  | 284<sub>id</sub> |  |  | 279<sub>id</sub> |  |  |
| Observations | 2609 |  |  | 2590 |  |  | 2554 |  |  |
| Marginal R<sup>2</sup> / Conditional R<sup>2</sup> | 0.010 / 0.521 |  |  | 0.185 / 0.521 |  |  | 0.272 / 0.601 |  |  |

Note. All continuous variables were standardized due to variables being on different scales. Prior to standardizing, PINES expression scores, age, education, trait negative affect, and trait positive affect were grand mean centered; daily positive affect was person-mean centered.

### Hourly negative affect

Detailed statistical results of mean hourly negative affect regressed on variables of interest are displayed in Table 4. In unadjusted models, PINES expression scores were not associated with hourly negative affect (*p* = .104). However, PINES expression scores were positively associated with hourly negative affect after controlling for demographic covariates and trait negative affect (*p* = .031, see Figure 2). The association between PINES expression scores and hourly negative affect remained similar after additionally controlling for trait positive affect and hourly positive affect (*p* = .031). Observed patterns remained unchanged when aggregating hourly data into a daily average (see supplemental material). No influential outliers were detected, and PINES expression scores remained a significant predictor of hourly negative affect after excluding data from practice days (*p* = .028). Lastly, these patterns were specific to negative affect, as PINES expression scores were not predictive of hourly positive affect (*p* = .289).

**Figure 2.**
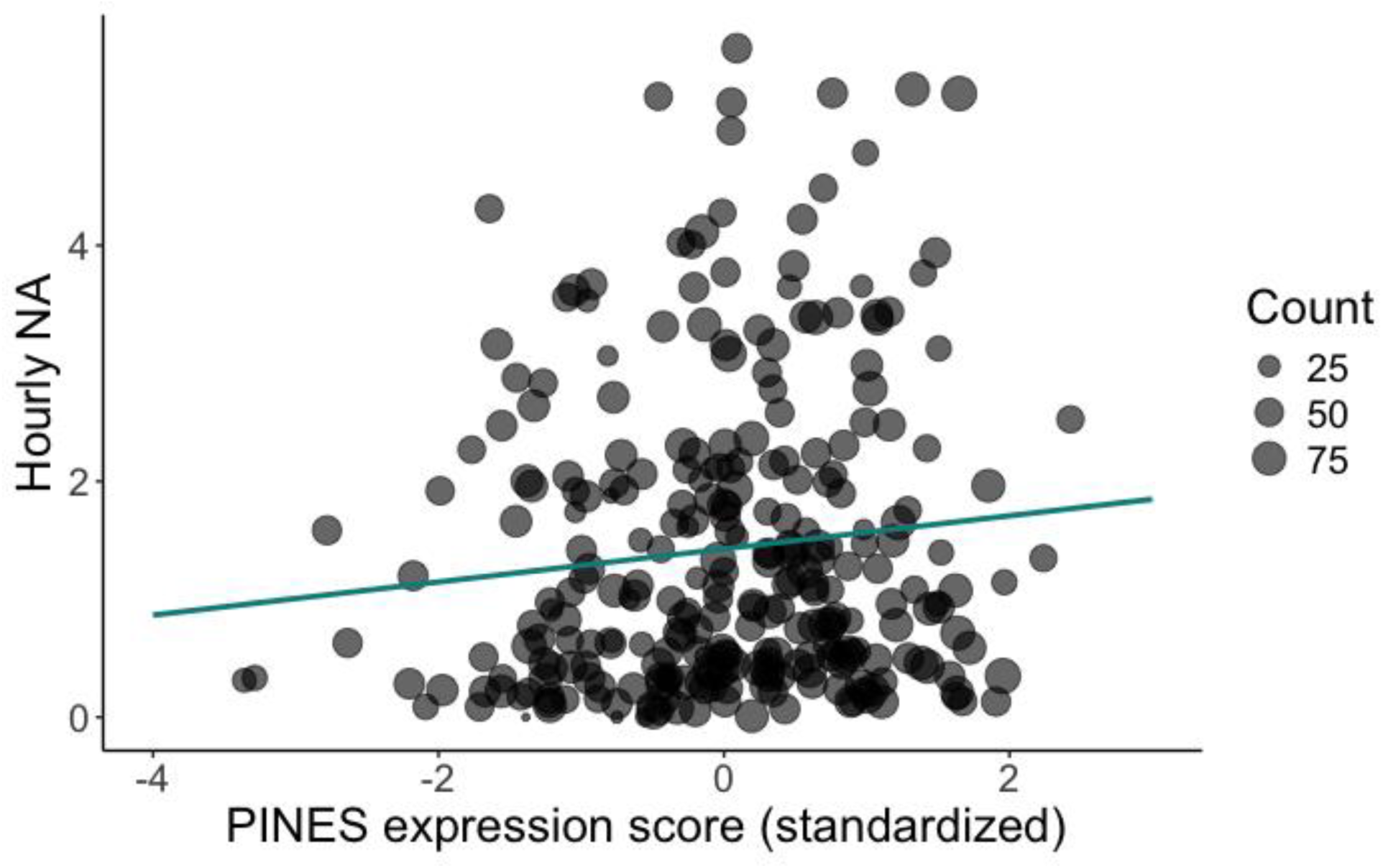
The association between PINES expression scores and hourly negative affect (NA). *Note:* This reflects the model that controlled for age, sex, education, and trait negative affect. For graphical purposes, individual data points were aggregated into a mean for each subject, such that each subject is represented by a dot and the size of the dot represents the number (count) of hourly entries across the entire observation period.

**Table 4.**
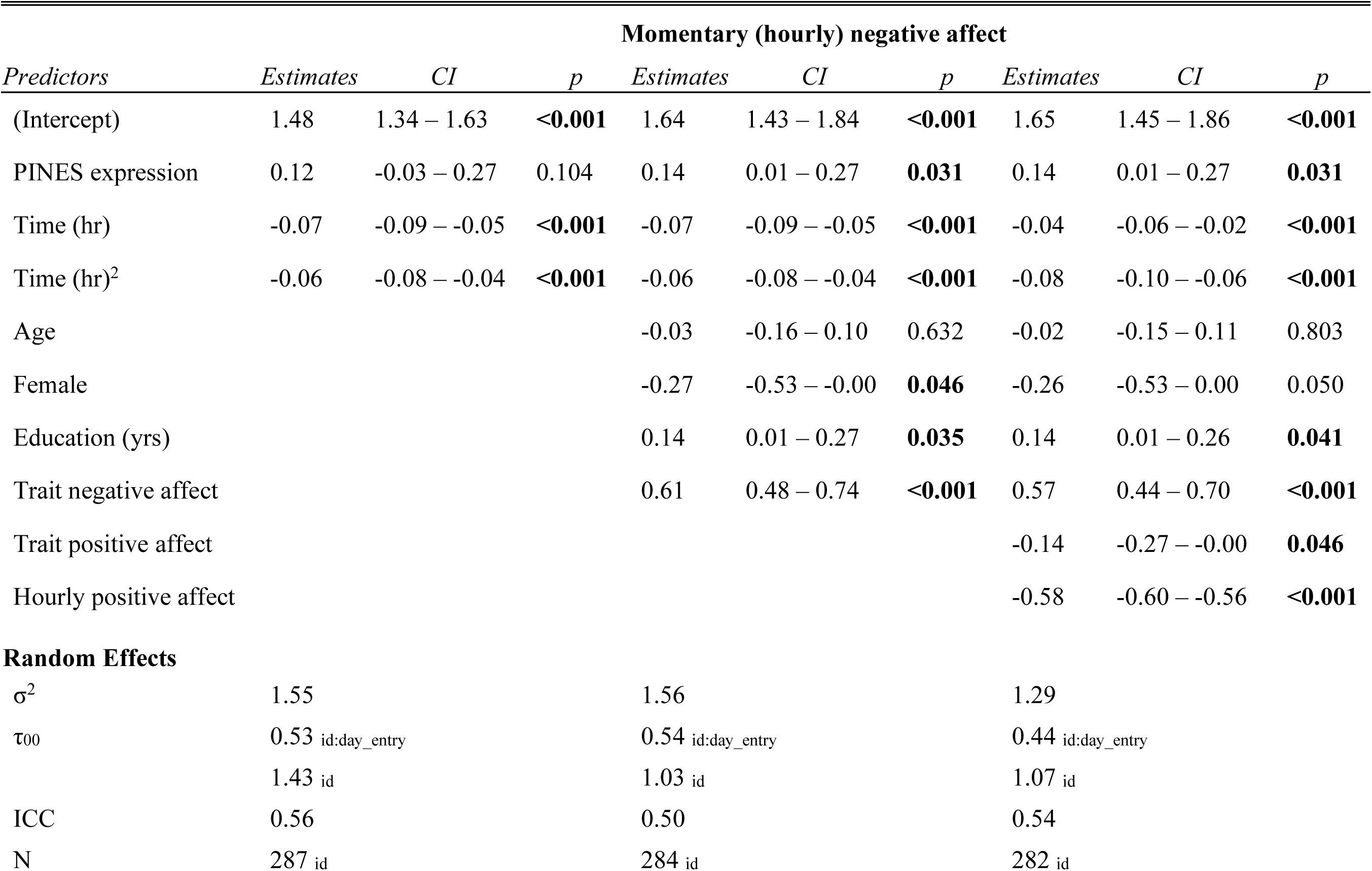
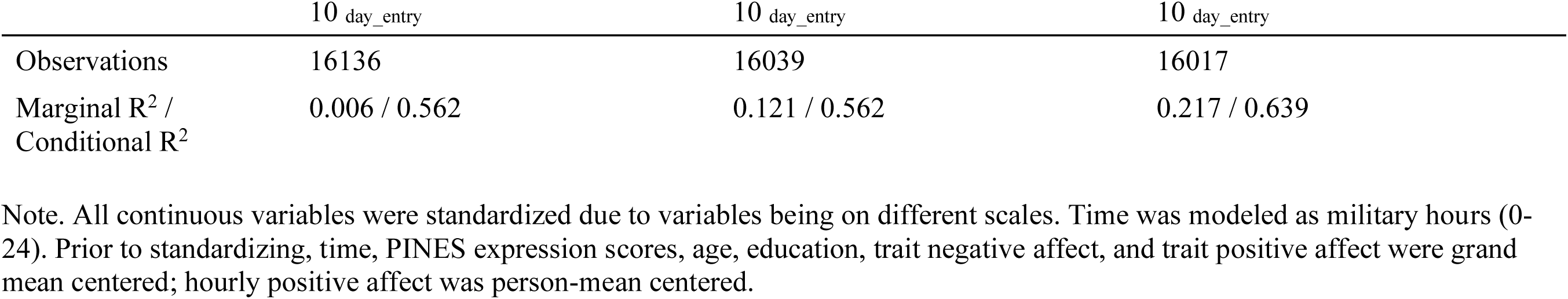
Linear mixed model results regressing hourly negative affect on PINES expression scores and covariates.

### Validity of the PINES and affective task

PINES significantly and positively predicted within-subject negative affect ratings, overall *rho* [95% CI] = .32 [.29 .40], *p* < .001; M±SD within-subject *r* = .51±.53, 50% subjects with *r* > .75; linear mixed effects model β(SE) = .32(.03), *t*(1058) = 12.28, p < .001 (see supplemental material for validation approach and results).

In analyses regressing everyday negative affect on task-induced negative affect ratings to Look Negative vs. Look Neutral images, higher task-induced negative affect was associated with higher daily *(p* = .008) and momentary negative affect *(p* = .025) after controlling for covariates (see supplemental material).

### Amygdalar contributions

Amygdala activity did not associate with daily (*p* = .235) or hourly negative affect (*p* = .662). PINES expression scores derived from the amygdala-lesioned PINES map significantly associated with daily (*p* = .008) and hourly negative affect (*p* = .030). Together, these findings suggest that amygdala activity was neither necessary nor sufficient to predict state negative affect in everyday life. Detailed statistics can be found in supplemental material.

## Discussion

Researchers have used fMRI and EMA to understand the relevance of task-evoked brain activity for predicting daily emotional and social behavior. However, as noted by a recent systematic review (Gadassi-Polack et al., 2024), existing studies are limited by small sample sizes (67% with less than 50 subjects) and a predominant focus on ROI analysis. The present study addresses the shortcomings of prior work by being one of the first to examine associations between a multivariate brain pattern of negative emotion and daily emotional patterns in a midlife sample of 287 subjects. We found that the degree of similarity to a whole-brain pattern of negative affect positively correlates with daily negative affect intensity. These patterns remained consistent regardless of how state negative affect was acquired: higher PINES expression scores were associated with higher end-of-day negative affect and higher hourly negative affect, after adjusting for covariates. By using a reproducible, brain signature of negative affect in conjunction with EMA, we provide the first evidence to our knowledge that variations in task-evoked brain patterns of negative affect reflect individual differences in state negative affect intensity in the real world, supporting the ecological validity of some multivariate brain patterns derived from task-fMRI.

Unlike self-reported hourly and daily negative affect, which were both associated with trait negative affect, PINES expression was only associated with daily and hourly negative affect, not trait negative affect. In other words, the PINES pattern appears to reflect state negative affect that is unconfounded by dispositional negative affect. Moreover, the PINES pattern was specific to negatively valenced emotion, as it was unrelated to daily and hourly positive affect. These findings provide evidence that some multivariate brain patterns, in this case the PINES, may have external validity: although the PINES was solely trained to predict state negative affect intensity in the scanner (Chang et al., 2015), PINES expression was associated with individual differences of the same psychological construct in a different context (i.e., state negative affect in daily life). These observations suggest that the PINES pattern captures a level of stability specific to negative affect experiences in acute situations. Said differently, while PINES expression cannot differentiate individuals who ascribe to more/less negative affect generally, PINES expression *can* differentiate individuals who tend to report more/less negative affect in time-restrained and, potentially, context-independent measurement periods. Thus, the PINES pattern may not be appropriate to use as a brain-based measure of negative affectivity – a stable personality trait characterized by persistent tendency to experience negative emotional states – and, instead, be better utilized as a brain-based measure of negative affective responding and experience. The latter findings agree with those of a recent multiverse study that failed to identify a reliable brain signature of neuroticism (Sicorello et al., in press).

Importantly, our findings corroborate pre-existing studies, which provided glimpses into the neurobiology of everyday experiences of negative affect. Indeed, select brain regions comprising the PINES pattern have separately been shown to correlate with negative affect in daily life. For example, task-induced functional activity in the anterior insula and the cingulate cortex – regions involved in interoception and emotional awareness and regulation – was associated with average daily negative affect and/or stress-induced negative affect in everyday life (Hur et al., 2022; Vaessen et al., 2023). Functional connectivity within the default mode network (DMN) – a network consisting of the posterior cingulate cortex, precuneus, medial prefrontal cortex – and connectivity between the insula and the DMN also correlated with daily negative affect in an adolescent and healthy adult sample (Ismaylova et al., 2018; Kaiser et al., 2019). Interestingly, associations between real-world experiences and subcortical regions typically implicated in affect and pain processing – such as the amygdala and periaqueductal gray – have been inconsistently observed, as threat-induced activity in these regions were linked with daily social stress (Eisenberger et al., 2007) but not daily negative affect (Hur et al., 2022). Together, these studies suggest that several regions likely play a role in daily emotion experiences, but isolated measures of brain activity may sometimes provide an incomplete story relative to multivariate patterns. Indeed, we also showed that amygdala activation failed to predict daily negative affect, nor was it necessary to the PINES’ prediction of daily negative affect.

This study has several strengths. By using a reproducible whole-brain pattern within a relatively large, healthy midlife sample, we increased the confidence, replicability, and generalizability of our findings. Moreover, we employed multiple techniques to measure negative affect in daily life. Capturing affective processes with (i.e., end-of-day reports) and without short periods of recall (i.e., hourly reports) minimized the influence of recall bias and added a form of internal replication to our findings. Finally, because the hourly and end-of-day scores were derived from only partially overlapping periods, we expanded the “cross section” of daily affective processes we examined by using both, providing a more reliable measure of an individual’s typical state affect.

Findings should be considered in light of some limitations. First, observed patterns may look different in a more representative sample, as our participants were predominantly White with a college education. Second, data collection occurred across several visits to minimize participant burden. Consequently, the first day of EMA data was collected, on average, 44 days before the fMRI scan. Although it was assumed that patterns would not change substantively within a span of weeks, the stability of multivariate brain patterns over a short period of time remains unclear. However, whether the fMRI scan preceded or followed EMA data collection does not alter the interpretation of our findings, which relied on correlational, not unidirectional, associations between the PINES pattern and daily negative affect. Third, although the PANAS is moderately stable over time (Watson & Walker, 1996), traditional measures of trait negative affect via the PANAS may partially reflect one’s semantic beliefs about themselves rather than a trait-like tendency to experience negative affect. Because one-time self-report and ambulatory self-report measures capture qualitatively different aspects of the conscious self (Conner & Barrett, 2012), future studies should consider alternative strategies to disentangle one’s belief from one’s actualized experience of trait-like negative affect. Lastly, because the PINES was trained to predict task-induced negative affect, the strength of the association between PINES expression and everyday negative affect may be limited by the ecological validity of the affective task itself. Our ancillary findings – namely that greater task-induced negative affect was associated with more daily and hourly negative affect in everyday life – were consistent with one prior study showing that greater negative bias toward laboratory stimuli (faces and scenes) was associated with heightened daily negative affect (Puccetti et al., 2023). The fact that the PINES and task-induced affect ratings correlated with each other *and* predicted everyday negative affect suggest that both measures capture overlapping aspects of state negative affect generalizable to everyday negative affect.

Despite widespread use of fMRI for understanding human emotion and behavior, robust support for the external validity of task-elicited brain activity for emotional experiences remains limited. The present study is among the first to demonstrate that individual differences in the expression of a multivariate brain pattern of negative affect corresponds to negative affect intensity in everyday life. Findings suggest that, despite the highly controlled and simplified nature of affect-related fMRI tasks, neural patterns derived from task-fMRI meaningfully relate to emotional patterns in the real world. Rigorous methodology performed in a relatively large sample, including validation of the PINES in a new cohort, strengthen the confidence and generalizability of our findings.

