## Supplemental Material for "From the scanner to daily life: Neural expression of the Picture-Induced Negative Emotion Signature is associated with real-world daily and momentary negative affect in midlife adults"

### Exclusion criteria for NOAH participants

NOAH participants were excluded if they were: diagnosed with cardiovascular or cerebrovascular disease, liver disease, lung disease, Type 1 diabetes, psychiatric or neurological disorder, kidney disease; had high resting blood pressure (≥160/100 mmHg) determined in the laboratory; received weight loss surgery or medication; consumed greater than 5 servings of alcohol per day; received cancer or chemotherapy treatment; were colorblind or had significant visual impairment; required an assistive walking device; were a shift worker; had barriers to completing electronic diary and ambulatory data collection (e.g., limited phone access); took antihypertensive, cardiac, anticonvulsant, anti-Parkinson, anti-HIV (e.g., protease inhibitors), prescription weight loss, immunosuppressant, chemotherapy, mania or centrally active psychotropic medications (excluding anti-anxiety, antidepressant, and antipsychotic/tranquilizer) on one or more occasions in the past 14 days; took anti-anxiety, anti-depressant, antipsychotic/tranquilizer, glucocorticoids (e.g., oral prednisone, cortisol), medical marijuana, sleep (e.g., trazadone), or asthma medications on a "regular" basis (defined as 7 or more days in the past 14 days); or took more than 2 non-insulin medication for diabetes on a "regular" basis. This is the first report of data from the emotion processing fMRI task and daily life data from the NOAH cohort.

#### IAPS IDs used

Unpleasant IAPS IDs used in the “Look Negative” condition were: 3350, 3120, 9414, 3051, 3550, 2053, 3500, 2703, 9400, 9040, 9921, 3102, 9050, 6831, 9252. Unpleasant IAPS IDs used in the “Regulate Negative” condition were: 9250, 9420, 3230, 3530, 6520, 3030, 3100, 9910, 2683, 3110, 6838, 6212, 9254, 9410, 3170. IAPS IDs used in the “Look Neutral” condition were: 7550, 2595, 2102, 2272, 2390, 2487, 2036, 2308, 2411, 2393, 9210,8312, 2026,7130, 2377.

### Validation of the PINES model in the NOAH cohort.

To our knowledge, no studies have yet tested the validity and generalizability of the PINES in a new cohort that underwent the same IAPS viewing task. Hence, we tested if PINES pattern expression scores were associated with in-scanner negative affect ratings in the present cohort.

For each subject, we estimated additional first-level General Linear Models (GLMs) in accordance with the approach specified by Figure 1B and text in Chang et al (2015). Specifically, subject-specific idiographic GLMs included up to 5 regressors reflecting Look-Negative and Look-Neutral image-viewing epochs that were modeled separately according to each rating level (1 to 5). As in Chang et al., a regressor reflecting all rating periods was included to account for motor activity, as well as a regressor reflecting the mean viewing epoch for all Look-Negative and Look-Neutral trials. Finally, we repeated the nuisance regression modeling of Chang et al., including 6 realignment parameters, their 1st derivatives, and the squares of these 12 regressors. As in Chang et al., we did not model Regulate-Negative trials for this validation analysis. Hence, idiographic subject-level GLMs produced up to 5 beta images per subject, depending on the diversity of a given subject's ratings, each reflecting image-viewing BOLD activity for that rating level. For each subject's rating-specific beta image, we computed the PINES pattern expression as the predicted rating and compared predictions to the corresponding observed rating.

Of the 295 participants with useable rating and imaging data, 68 (23%) gave a rating of 1 in one or more trials, 202 (69%) gave a rating of 2, 280 (96%) gave a rating of 3, 281 (96%) a rating of 4, and 229 (78%) a rating of 5. As for diversity of responses, one participant gave only 1 unique rating level, and were removed from further analysis; 18 (6%) gave 2 unique rating levels; 105 (36%) gave 3 unique rating levels, 137 (47%) gave 4 unique rating levels, and 32 (11%) gave the full range of 5 rating levels.

Results showed that predicted (pattern expression) and observed ratings were positively associated across ratings and participants, Spearman's $rho = .34$, $p < .001$, 95% bootstrapped confidence intervals (CI) .29 to .40, average within-person predicted-observed Pearson correlation (for participants with 3+ unique rating levels, N=274) of $r \pm SD = .51 \pm .53$, with 50% of participants showing $r > .75$. A linear mixed effects model regressing pattern expressions on observed rating levels with subject as random effects also showed a positive association, $\beta(SE) = .32(.03)$, $t(1058) = 12.28$, $p < .001$.

To summarize, applying the PINES model to idiographic ratings-specific images in the present cohort generated significant positive predictions of observed negative affect ratings. This indicates that pattern expression scores elicited by the PINES are statistically associated with in-scanner negative affect ratings in the present cohort, owing to the validity and generalizability of the PINES in the broader context of our findings bearing on daily-life negative affect.

**Table S1.** Linear mixed model results regressing hourly negative affect on PINES expression and covariates, excluding data from practice days

| | **Hourly NA** | | |
|---|---|---|---|
| *Predictors* | *Estimates* | *CI* | *p* |
| (Intercept) | 1.70 | 1.48 – 1.91 | **<0.001** |
| PINES expression | 0.15 | 0.02 – 0.28 | **0.028** |
| Time (hr) | -0.06 | -0.08 – -0.04 | **<0.001** |
| Time $(hr)^2$ | -0.06 | -0.09 – -0.04 | **<0.001** |
| Age | -0.03 | -0.17 – 0.11 | 0.671 |
| Female | -0.31 | -0.58 – -0.03 | **0.030** |
| Education (yrs) | 0.18 | 0.04 – 0.31 | **0.009** |
| Trait negative affect | 0.60 | 0.47 – 0.73 | **<0.001** |
| **Random Effects** | | | |
| $\sigma^2$ | 1.56 | | |
| $\tau_{00\ id:day\_entry}$ | 0.44 | | |
| $\tau_{00\ id}$ | 1.14 | | |
| ICC | 0.50 | | |
| $N_{id}$ | 284 | | |
| $N_{day\_entry}$ | 10 | | |
| Observations | 14188 | | |
| Marginal $R^2$ / Conditional $R^2$ | 0.122 / 0.565 | | |

Note. All continuous variables were standardized due to variables being on different scales. Time was modeled as military hours (0-24). Prior to standardizing, time, PINES expression scores, age, education, and trait negative affect were grand mean centered.

**Table S2.** Linear mixed model results regressing aggregated hourly negative affect on PINES expression and covariates.

| | Daily aggregated hourly negative affect | | | | | | | | |
|---|---|---|---|---|---|---|---|---|---|
| *Predictors* | *Estimates* | *CI* | *p* | *Estimates* | *CI* | *p* | *Estimates* | *CI* | *p* |
| (Intercept) | 1.45 | 1.31 – 1.59 | **<0.001** | 1.59 | 1.39 – 1.79 | **<0.001** | 1.59 | 1.39 – 1.79 | **<0.001** |
| PINES expression | 0.12 | -0.02 – 0.26 | 0.104 | 0.13 | 0.01 – 0.26 | **0.036** | 0.13 | 0.01 – 0.26 | **0.037** |
| Age | | | | -0.02 | -0.15 – 0.10 | 0.712 | -0.01 | -0.14 – 0.11 | 0.828 |
| Female | | | | -0.24 | -0.50 – 0.02 | 0.065 | -0.23 | -0.49 – 0.03 | 0.082 |
| Education (yrs) | | | | 0.12 | -0.01 – 0.24 | 0.066 | 0.11 | -0.01 – 0.24 | 0.074 |
| Trait negative affect | | | | 0.63 | 0.50 – 0.75 | **<0.001** | 0.59 | 0.46 – 0.72 | **<0.001** |
| Trait positive affect | | | | | | | -0.12 | -0.25 – 0.01 | 0.076 |
| Daily positive affect | | | | | | | -0.33 | -0.38 – -0.28 | **<0.001** |
| **Random Effects** | | | | | | | | | |
| $\sigma^2$ | 0.94 | | | 0.95 | | | 0.84 | | |
| $\tau_{00}$ | 1.35 $_{id}$ | | | 0.93 $_{id}$ | | | 0.95 $_{id}$ | | |
| ICC | 0.59 | | | 0.50 | | | 0.53 | | |
| N | 287 $_{id}$ | | | 284 $_{id}$ | | | 282 $_{id}$ | | |
| Observations | 1539 | | | 1528 | | | 1522 | | |
| Marginal $R^2$ / Conditional $R^2$ | 0.006 / 0.592 | | | 0.189 / 0.592 | | | 0.229 / 0.638 | | |

Note. All continuous variables were standardized due to variables being on different scales. Time was modeled as military hours (0-24). Prior to standardizing, time, PINES expression scores, age, education, trait negative affect, and trait positive affect were grand mean centered; daily positive affect was person-mean centered

**Table S3.** End-of-day negative affect regressed on mean amygdala activity (Look Negative vs. Look Neutral) and covariates

| | End-of-day negative affect | | |
|---|---|---|---|
| *Predictors* | *Estimates* | *CI* | *p* |
| (Intercept) | 2.02 | 1.77 – 2.27 | **<0.001** |
| Amygdala activity | 0.09 | -0.06 – 0.25 | 0.234 |
| Age | -0.11 | -0.27 – 0.05 | 0.160 |
| Female | 0.06 | -0.27 – 0.38 | 0.731 |
| Education (yrs) | 0.17 | 0.02 – 0.33 | **0.031** |
| Trait negative affect | 0.85 | 0.69 – 1.01 | **<0.001** |
| **Random Effects** | | | |
| $\sigma^2$ | 2.14 | | |
| $\tau_{00\ id}$ | 1.53 | | |
| ICC | 0.42 | | |
| $N_{id}$ | 284 | | |
| Observations | 2590 | | |
| Marginal $R^2$ / Conditional $R^2$ | 0.178 / 0.521 | | |

Note. All continuous variables were standardized due to variables being on different scales. Prior to standardizing, time, PINES expression scores, age, education and trait negative affect.

**Table S4.** Hourly negative affect regressed on mean amygdala activity (Look Negative vs. Look Neutral) and covariates

| | Hourly negative affect | | |
|---|---|---|---|
| *Predictors* | *Estimates* | *CI* | *p* |
| (Intercept) | 1.63 | 1.42 – 1.83 | **<0.001** |
| Amygdala activity | 0.03 | -0.10 – 0.16 | 0.661 |
| Time (hr) | -0.07 | -0.09 – -0.05 | **<0.001** |
| Time (hr)$^2$ | -0.06 | -0.08 – -0.04 | **<0.001** |
| Age | -0.03 | -0.16 – 0.10 | 0.657 |
| Female | -0.24 | -0.50 – 0.02 | 0.074 |
| Education | 0.14 | 0.01 – 0.27 | **0.041** |
| Trait negative affect | 0.61 | 0.48 – 0.74 | **<0.001** |
| **Random Effects** | | | |
| $\sigma^2$ | 1.56 | | |
| $\tau_{00}$ id:day_entry | 0.54 | | |
| $\tau_{00}$ id | 1.05 | | |
| ICC | 0.50 | | |
| N id | 284 | | |
| N day_entry | 10 | | |
| Observations | 16039 | | |
| Marginal $R^2$ / Conditional $R^2$ | 0.116 / 0.562 | | |

Note. All continuous variables were standardized due to variables being on different scales. Time was modeled as military hours (0-24). Prior to standardizing, time, PINES expression scores, age, education and trait negative affect.

**Table S5.** End-of-day negative affect regressed on amygdala-lesioned PINES pattern expression and covariates

| | End-of-day negative affect | | |
|---|---|---|---|
| *Predictors* | *Estimates* | *CI* | *p* |
| (Intercept) | 1.45 | 0.96 – 1.93 | **<0.001** |
| PINES expression (w/o amygdala) | 9.49 | 2.49 – 16.49 | **0.008** |
| Age | -0.12 | -0.28 – 0.04 | 0.141 |
| Female | 0.02 | -0.30 – 0.34 | 0.895 |
| Education (yrs) | 0.17 | 0.02 – 0.33 | **0.030** |
| Trait negative affect | 0.85 | 0.70 – 1.01 | **<0.001** |
| **Random Effects** | | | |
| $\sigma^2$ | 2.14 | | |
| $\tau_{00\ id}$ | 1.50 | | |
| ICC | 0.41 | | |
| $N_{id}$ | 284 | | |
| Observations | 2590 | | |
| Marginal $R^2$ / Conditional $R^2$ | 0.185 / 0.521 | | |

Note. All continuous variables were standardized due to variables being on different scales. Prior to standardizing, time, PINES expression scores, age, education and trait negative affect.

**Table S6.** Hourly negative affect regressed on amygdala-lesioned PINES pattern expression and covariates

| | Hourly negative affect | | |
|---|---|---|---|
| *Predictors* | *Estimates* | *CI* | *p* |
| (Intercept) | 1.64 | 1.43 – 1.84 | **<0.001** |
| PINES expression (w/o amygdala) | 0.14 | 0.01 – 0.27 | **0.030** |
| Time | -0.07 | -0.09 – -0.05 | **<0.001** |
| Time (hr)$^2$ | -0.06 | -0.08 – -0.04 | **<0.001** |
| Age | -0.03 | -0.16 – 0.10 | 0.633 |
| Female | -0.27 | -0.53 – -0.00 | **0.047** |
| Education | 0.14 | 0.01 – 0.27 | **0.036** |
| Trait negative affect | 0.61 | 0.48 – 0.74 | **<0.001** |
| **Random Effects** | | | |
| $\sigma^2$ | 1.56 | | |
| $\tau_{00\ id:day\_entry}$ | 0.54 | | |
| $\tau_{00\ id}$ | 1.03 | | |
| ICC | 0.50 | | |
| $N_{id}$ | 284 | | |
| $N_{day\_entry}$ | 10 | | |
| Observations | 16039 | | |
| Marginal $R^2$ / Conditional $R^2$ | 0.121 / 0.562 | | |

Note. All continuous variables were standardized due to variables being on different scales. Time was modeled as military hours (0-24). Prior to standardizing, time, PINES expression scores, age, education and trait negative affect.

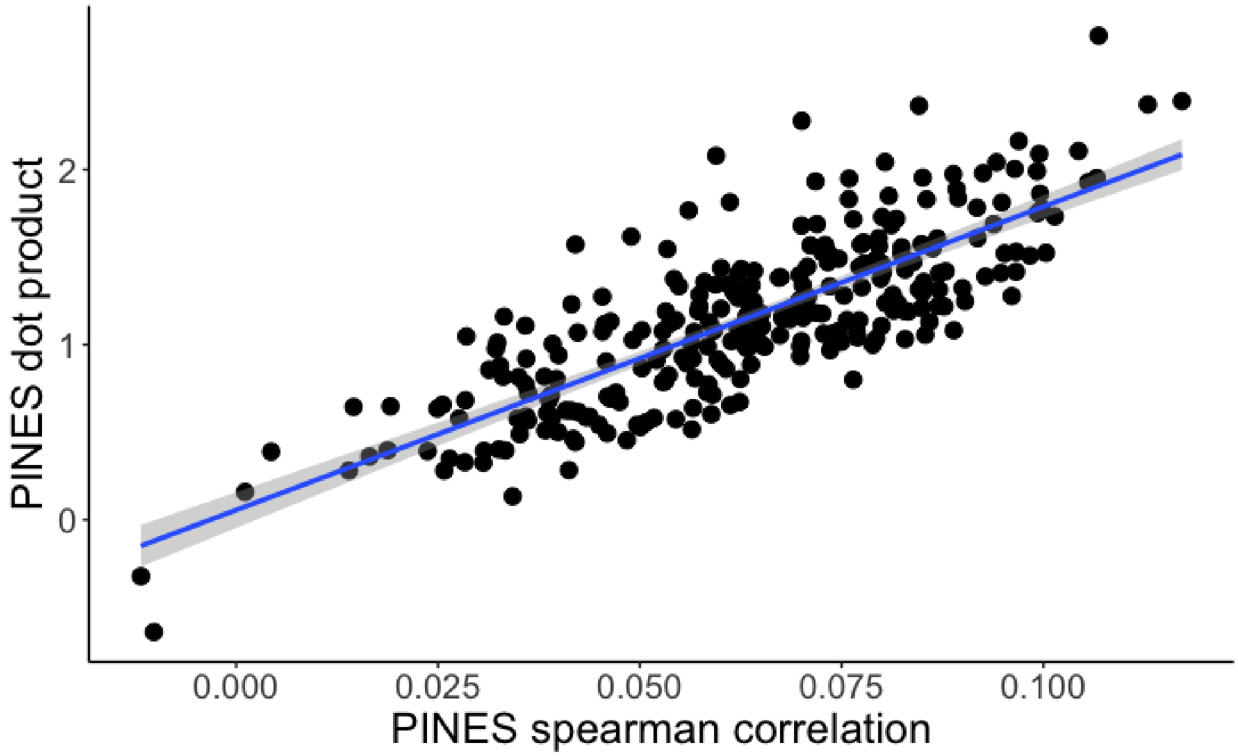

**Figure S1.** Scatterplot depicting correlation of PINES expression derived from Spearman's correlation and PINES expression derived from dotproduct (*r* = .80)

**Table S7.** End-of-day negative affect regressed on PINES expression derived from dot product and covariates

| | End-of-day negative affect | | |
|---|---|---|---|
| *Predictors* | *Estimates* | *CI* | *p* |
| (Intercept) | 2.04 | 1.79 – 2.29 | **< .001** |
| PINES expression (dot product) | 0.14 | -0.01 – 0.30 | 0.073 |
| Age | -0.10 | -0.26 – 0.06 | 0.205 |
| Female | 0.03 | -0.30 – 0.35 | 0.869 |
| Education (yrs) | 0.17 | 0.02 – 0.33 | **0.030** |
| Trait negative affect | 0.86 | 0.70 – 1.02 | **<0.001** |
| **Random Effects** | | | |
| $\sigma^2$ | 2.14 | | |
| $\tau_{00\ id}$ | 1.52 | | |
| ICC | 0.42 | | |
| N id | 284 | | |
| Observations | 2590 | | |
| Marginal $R^2$ / Conditional $R^2$ | 0.180 / 0.521 | | |

Note. All continuous variables were standardized due to variables being on different scales. Prior to standardizing, time, PINES expression scores, age, education and trait negative affect.

**Table S8.** Hourly negative affect regressed on PINES expression derived from dot product and covariates

| | Hourly negative affect | | |
|---|---|---|---|
| *Predictors* | *Estimates* | *CI* | *p* |
| (Intercept) | 1.64 | 1.43 – 1.84 | **<0.001** |
| PINES expression (dot product) | 0.10 | -0.03 – 0.22 | 0.145 |
| Time (hr) | -0.07 | -0.09 – -0.05 | **<0.001** |
| Time $(hr)^2$ | -0.06 | -0.08 – -0.04 | **<0.001** |
| Age | -0.02 | -0.15 – 0.11 | 0.748 |
| Female | -0.26 | -0.53 – 0.00 | 0.052 |
| Education | 0.14 | 0.01 – 0.27 | **0.036** |
| Trait negative affect | 0.61 | 0.49 – 0.74 | **<0.001** |
| **Random Effects** | | | |
| $\sigma^2$ | 1.56 | | |
| $\tau_{00\ id:day\_entry}$ | 0.54 | | |
| $\tau_{00\ id}$ | 1.04 | | |
| ICC | 0.50 | | |
| $N_{id}$ | 284 | | |
| $N_{day\_entry}$ | 10 | | |
| Observations | 16039 | | |
| Marginal $R^2$ / Conditional $R^2$ | 0.119 / 0.562 | | |

Note. All continuous variables were standardized due to variables being on different scales. Time was modeled as military hours (0-24). Prior to standardizing, time, PINES expression scores, age, education and trait negative affect.
